# Quantifying the Inhibitory Impact of Soluble Phenolics on Carbon Mineralization from *Sphagnum*-rich Peatlands

**DOI:** 10.1101/2021.05.24.445415

**Authors:** Alexandra B. Cory, Jeffrey P. Chanton, Robert G.M. Spencer, Virginia I. Rich, Carmody K. McCalley, IsoGenie Project Coordinators, Rachel M. Wilson, Scott R. Saleska, Patrick M. Crill, Gene W. Tyson, Ruth K. Varner, Matthew B. Sullivan, Steven Frolk

**Author notes:** IsoGenie Project Coordinators list of authors and affiliations appears in the supplement.

## Abstract

The mechanisms controlling the extraordinarily slow carbon (C) mineralization rates characteristic of *Sphagnum*-rich peatlands (“bogs”) remain somewhat elusive, despite decades of research on this topic. Soluble phenolic compounds have been invoked as potentially significant contributors to bog peat recalcitrance due to their affinity to slow microbial metabolism and cell growth. Despite this potentially significant role, the effects of soluble phenolic compounds on bog peat C mineralization remain unclear.

We analyzed this effect by manipulating the concentration of free soluble phenolics in anaerobic bog peat incubations using water-soluble polyvinylpyrrolidone (PVP), a compound that binds with and inactivates phenolics, preventing phenolic-enzyme interactions. CO_2_ and CH_4_ production rates (end-products of C mineralization) correlated positively with PVP concentration following Michaelis-Menten (M.M.) saturation functions. Using M.M. parameters, we determined that soluble phenolics inhibit, at minimum, 57 ± 16% of total C (CO_2_+CH_4_) mineralization in the anaerobic incubation conditions studied. These findings are consistent with other studies that have indicated that soluble phenolics play a significant role in regulating bog peat stability in the face of decomposition.

## Introduction

Due to the enormous quantity of carbon (C) contained in peatlands— current estimates ranging from ~530-1,175 Pg globally (equivalent to ~60% - 134% of current atmospheric C stores) [1–3]—shifts in peatland C cycling have potentially significant impacts on the global climate. Most peatlands are CO_2_ sinks and CH_4_ sources [4–6]. The former process is cooling to the climate, while the latter has a warming effect. For the past 8000-11,000 years, peatland C deposition has had a net cooling effect on the climate [7]. Over the last ~150 years, the peatland carbon sink has diminished with estimates of present climatic impacts ranging from slightly cooling (−0.7 W x m^−2^; instantaneous box-model estimate) [7] to slightly warming (+0.6 Pg CO_2_-equiv y^−1^; field flux estimate) [5].

This regime shift is the result of climate change-induced disruptions to the peatland system including water table shifts, permafrost thaw, and increased frequency of fire and drought [8–10]. Acute anthropogenic disturbances (e.g., drainage and burning) have also created significant C balance disruptions and will likely continue to do so without political intervention [5]. Together, these disruptions may speed up rates of CO_2_ and CH_4_ production via decomposition, thereby shifting peatlands into significant drivers of warming. It is thus imperative that we incorporate an accurate assessment of peatland-climate dynamics into global climate models. To do so, we must understand the underlying biogeochemical processes responsible for peat C mineralization.

Decomposition in *Sphagnum*-rich peatlands (“bogs”) is extraordinarily slow and the cause of this remains elusive despite decades of research [11–17]. Typically present in northern permafrost zones, bogs are saturated with cold, moderately acidic (pH~4.5) water introduced via gradual permafrost thaw [18]. The resulting anoxia precludes oxic respiration while the generally nutrient-deplete conditions characteristic of bogs hinder respiration via inorganic terminal electron acceptors (“TEAs”) [18]. As such, organic matter decomposition is limited primarily to three energy-limiting processes (1) hydrolysis (breakdown of complex organic compounds into simple compounds), (2) fermentation, and (3) methanogenesis. Though these conditions contribute to slow decomposition rates, they do not fully account for observed bog C mineralization rates.

Soluble phenolics have been invoked as potentially significant inhibitors of bog decomposition due to their propensity to suppress microbial metabolism and inhibit cell growth [19–24]. Metabolic disruptions are attributed to phenolic-enzyme interactions (bonding and/or adsorption), which limit enzyme activity [25–30]. Disruptions to cell growth and function are attributed to phenolic-membrane interactions, which can cause membrane injury and increased permeability. The latter is associated with increased influx of extracellular compounds—some of which can be toxic to micro-organisms—and increased efflux of intracellular components that are necessary for cell growth, such as proteins, potassium, and phosphates [31–35].

Though the inhibitory effects of soluble phenolics are generally accepted, the extent to which they inhibit C mineralization in *Sphagnum*-rich peatlands (“bogs”) remains unclear [9,20,21,36,37]. Studies to date have focused heavily on the potential for soluble phenolics to inhibit enzymatic hydrolysis. These studies have yielded conflicting assessments of this role, ranging from significant [9,20,21] to inconsequential [36,37]. To clarify the impacts of soluble phenolics on bog C mineralization, it is necessary to incorporate all three stages of bog decomposition (hydrolysis, fermentation, and methanogenesis) into this assessment. There are three reasons for this need: (1) inhibition of fermentation and methanogenesis by soluble phenolics has been observed [24,31,38,39]; (2) evidence of simple sugar buildup in bog peat indicates that C mineralization rates are sometimes not limited by hydrolysis (which produces simple sugars), but rather fermentation and/or methanogenesis (which collectively consume simple sugars) [40]; and (3) the relationship between respiration rates and hydrolytic enzyme activities is sometimes shown to be insignificant [41].

Studies that consider the impacts of soluble phenolics on all three stages of bog decomposition are scant. Suppression of CO_2_ and CH_4_ production—end-products to all three stages—has been observed in incubated peat amended with phenolic-rich dissolved organic matter (“DOM”) [42]. Though it is feasible that soluble phenolics caused this inhibition, the presence of other potentially inhibitory DOM compounds precludes definitive affirmation of this effect [42]. Suppression of aerobic respiration by soluble phenolics has been observed in an aerobic incubation experiment [9], but only after the addition of respiration substrates (C and nutrients). This finding suggests that the potentially inhibitory impacts of soluble phenolics are inconsequential in substrate-limited settings. Given that substrate supply varies in response to site-specific factors—such as vegetation and climate—it is necessary to broaden assessments regarding the impact of soluble phenolics on C mineralization to more sites. Moreover, given that most bog peat typically resides in anaerobic conditions [18], it is necessary to apply these assessments to anaerobic conditions.

We analyzed the relationship between soluble phenolic content and anaerobic C mineralization rates in a bog site within a Swedish permafrost peatland (Stordalen Mire). We used the cumulative production of CO_2_ and CH_4_ (the end-products of C mineralization in anaerobic bog environments) to determine C mineralization rates. Using the methods of [43], we manipulated the concentration of free soluble phenolics using water-soluble polyvinylpyrrolidone (PVP). This synthetic polymer “inactivates” soluble phenolics via binding and precipitation, preventing phenolic-enzyme interactions from occurring [44]. To maximize this inactivation, we sought to saturate our incubations with PVP. As the concentration necessary to achieve saturation in our incubations was unknown, additions were undertaken across a wide concentration range.

We hypothesized that increasing PVP concentration would increase C mineralization rates without the addition of respiration substrates because substantial buildup of simple sugars has been observed at our study site [40]. We hypothesized that the relationship between PVP concentration and C mineralization rates would follow a Michaelis-Menten saturation function (Fig 1). By examining the effects of PVP saturation on C mineralization, we sought to identify the extent to which soluble phenolics inhibit bog C mineralization. We expect this to be a minimum estimate given that (1) even in PVP-saturated conditions, a minute portion of free soluble phenolics likely persist [44] and (2) of these persisting soluble phenolics, some could feasibly continue interacting with enzymes, leading to continued inhibition of C mineralization.

**Fig 1.**
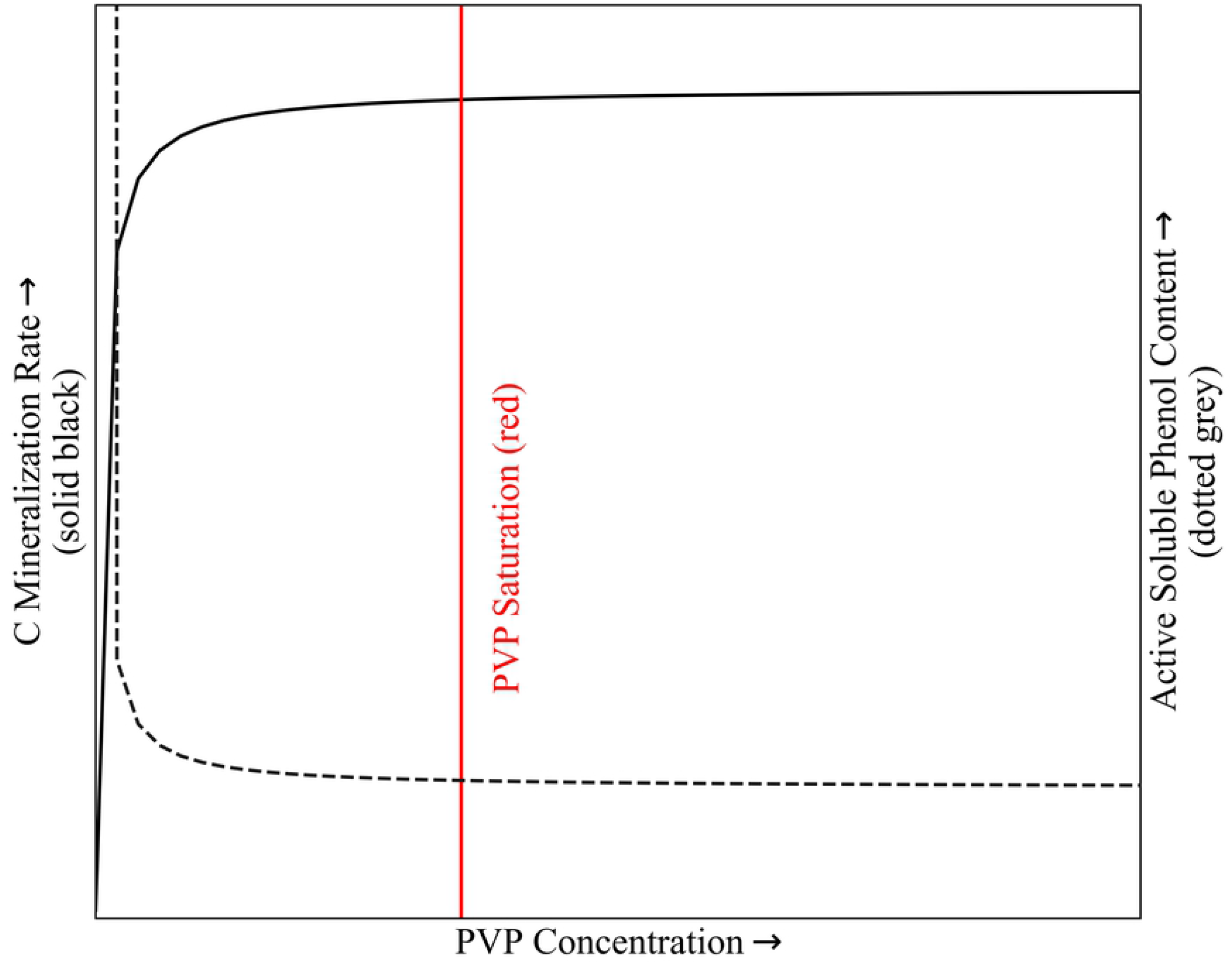
Hypothesized relationship between polyvinylpyrrolidone (PVP) concentration vs. C mineralization. C mineralization rate (measured by CO_2_ and CH_4_ production) corresponds to the primary y axis (solid black line). Assumed active soluble phenol content corresponds to the secondary y axis (dotted grey line). Addition of PVP was hypothesized to positively impact CO_2_ and CH_4_ production rates by inactivating soluble phenolics which would otherwise inhibit C decomposition. This relationship was expected to follow a Michaelis-Menten saturation curve. After reaching a point of PVP saturation (red line), further increases in PVP concentration were expected to yield no significant changes in C mineralization rates.

### Site Description

Stordalen Mire is a subarctic peatland approximately 10 km east of Abisko, Sweden (68.35°, 19.05°E). Located within the discontinuous permafrost zone, it is comprised of three dominant habitats—palsas, bogs, and fens—reflecting various stages of thaw with different hydrologic regimes. For an in-depth description of these habitats, see [45–47]. Peat samples for this study were collected from a bog site within the Mire, which falls mid-stage along the permafrost thaw progression. The bog site is perched above the regional water table separated by a layer of permafrost. As such, its water inputs are limited to rainfall (“ombrotrophic”). Vegetation is dominated by mosses (mainly *Sphagnum* spp.), with some sedges (e.g. *Eriophorum vaginatum, Carex rotundata*).

## Methods

### Field Collection

Peat for the laboratory incubation experiments was obtained in July 2018 using an Eijenkamp perforated stainless steel corer. We sectioned the core using a razor blade and set aside peat from 9-19 cm depth for incubation analysis. The peat was stored at −20°C from the point of collection up until the experiment start date.

### Experimental Design

We prepared six treatments, each with three replicates, totaling 21 vials. Treatment 1 was an untreated control. Treatments 2-6 spanned a polyvinylpyrrolidone (PVP) concentration range of 0.001-0.064 g/mL (Table 1), which extended above and below concentrations shown to stimulate enzymatic hydrolysis in DOM-rich arctic river water (0.005 and 0.010 g/mL) [43].

**Table 1.** PVP Concentration by treatment number.

| Treatment # | PVP Conc. (g/mL) |
| --- | --- |
| 1 | 0 |
| 2 | 0.001 |
| 3 | 0.004 |
| 4 | 0.016 |
| 5 | 0.032 |
| 6 | 0.064 |

Each treatment was conducted in triplicate, totaling 21 vials. Each vial consisted of saturated shallow (9-19 cm) bog peat. Vials were held in dark, anoxic conditions.

We thawed and homogenized the peat by de-clumping it with gloved hands and forceps. We aliquoted 40 g of homogenized peat and 30 mL deionized (DI) water into each 160-mL clear borosilicate vial. We then capped the vials with rubber septa and sealed them with aluminum crimps. To create anoxic conditions, we vigorously shook the vials and flushed the headspace with N_2_ gas. We repeated this process until headspace CO_2_ concentrations measured less than 0.1% (see “Gas Analysis”), which was two magnitudes lower than the CO_2_ production measured during the experiment. The above-gauge headspace pressure was ~3.5 psi immediately following headspace flushing. We stored the vials in total darkness at room temperature (20-22°C) and allowed them to sit for a 25-day pre-incubation phase. The purpose of this step was to facilitate consumption of molecular O2 and reduction of oxidized chemical species [48,49]. We reference all time points relative to the end of this pre-incubation period such that day 1 corresponds to the first day following the 25-day pre-incubation phase.

On day 1, water-soluble polyvinylpyrrolidone PVP was added (Sigma Aldrich, CAS #: 9003-39-8, average molecular weight: 40,000). Using a stock solution of PVP at 0.256 g/mL in DI water, we performed serial dilutions to obtain the final PVP concentrations detailed in Table 1, except for the control (treatment 1), which was composed entirely of DI water. We injected 10 mL of the relevant PVP solution (except for Treatment 1, which contained only DI water) into each vial to achieve the final concentrations indicated in Table 1. The final ratio of grams wet peat: mL solution was 1:1, which was enough to fully saturate the peat.

After addition of the PVP solution, we re-flushed the headspace with N2 gas and shook the vials until we once again measured headspace CO_2_ concentrations less than 0.1%. At this time, the above-gauge headspace pressure was once again ~3.5 psi. We periodically measured headspace pressure to ensure that it did not fall below 0.5 psi (to prevent air infiltration). Since the volume extracted for gas analysis (≤ 250 μl) was a small fraction of the total headspace volume (80 mL), gas replacement was not necessary to maintain >0.5 psi headspace pressures. We collected headspace samples for analysis of CO_2_ and CH_4_ concentrations every 3-10 days throughout the duration of the experiment (56 days; consistent with [48]). At the end of the experiment, we removed the vial caps and dried the samples at a constant 68°C. Once the sample weights stabilized, we obtained final dry weights, which we used to calculate CO_2_ and CH_4_ production per g dry peat (see “Statistical Analysis”).

### Gas Analysis

We performed all CO_2_ and CH_4_ concentration analyses via Flame-Ionization-Detector Gas Chromatography (GC-FID) using methods established by [46]. We used a gas-tight syringe for injection of all samples and standards. The GC flow rate was 30 mL/min, and the temperatures were 140, 160, and 380°C for the column, detector, and methanizer, respectively. On each sampling day, we created a linear calibration curve for both CO_2_ and CH_4_ using standards of known concentrations. Before sample analyses, we shook the vials vigorously to liberate gases trapped in the peat pore-spaces. We also recorded headspace pressures to calculate partial pressures of CO_2_ and CH_4_ (which were essential for statistical analysis).

### Statistical Analysis

We calculated average production rates (μmoles × g dry peat^−1^ × day^−1^) for CO_2_ and CH_4_ using the steps outlined below. We determined total C mineralization rates (“C_tot_”) by taking the sum of CO_2_ and CH_4_ production rates.

To calculate gas production rates, we first determined the quantity of gas injected into the GC—n_gas(inj)_—by inputting sample peak amplitudes into our standard calibration curve (eqn. 1). We used the ideal gas law to determine the total moles injected—n_tot(inj)_ (eqn. 2). We calculated the gas fraction—F_gas_—using eqn. 3, which we used to calculate headspace partial pressures— P_gas_ (eqn. 4). We then applied this value to the ideal gas law to quantify the headspace moles— n_gas(HS)_ (eqn. 5). Using Henry’s Law (eqn. 6), we calculated the dissolved gas concentration— C_gas(aq.)_—which we used to determine the moles in the aqueous phase—n_gas_(aq) (eqn. 7). We determined the moles per vial—n_gas_ × vial^−1^—using eqn. 8. Our final daily production values— n_gas_ × g^−1^ —were obtained by eqn. 9 (where g=dry peat weight).

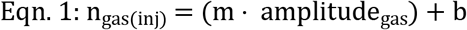

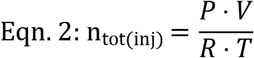

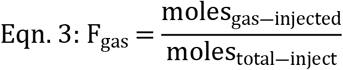

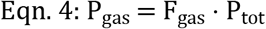

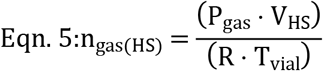

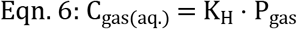

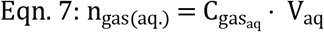

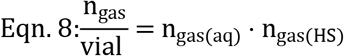

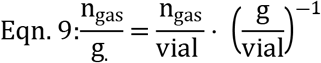

Production rate time series were best approximated using linear regression equations. We determined average production rates (n_gas_ × g^−1^ × d^−1^) using the slopes of these equations. We calculated respective R^2^ values using the Microsoft Excel RSQ function. To assess the significance of PVP concentration on production rates, we used the Microsoft Excel 2-tailed T-test (T.test) function.

### Modeled Response to PVP Addition

The relationship between gas production rate and PVP concentration was best approximated using Michaelis-Menten equations. Since production rates were expected to be nonzero in controls (where PVP=0 g × mL^−1^) we appended a y-intercept to the Michaelis-Menten equation (resulting in eqn. 10). The y-intercept—Prod_0_—was equivalent to the average production rate of controls (n=3).

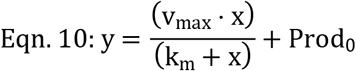

We used the curve_fit function from the Optimize package within Python’s SciPy library to determine the values of Michaelis-Menten constants v_max_ and k_m_. We determined the production rates in PVP-saturated peat—Prod_sat_—by summing the y-intercept (Prod_0_) and vmax using eqn. (11).

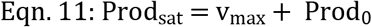

We used non-parametric bootstrapping (1,000 simulations) to calculate the standard deviation (95% confidence interval) for calculated v_max_ and k_m_ values. In addition to the Optimize package (referenced above), these simulations were programmed using the Numpy and Pandas Python data packages. To determine the fraction of observed variance explained by the Michaelis-Menten model, we calculated the R^2^ value for measured vs. modeled production rates.

## Results

When production rates of all three of CO_2_, CH_4_, and C_tot_ (CO_2_+CH_4_) follow similar trends, we will refer to them collectively as GHG_c_ production. GHG_c_ production rates increased with PVP concentration across the entire PVP concentration range studied (Fig 2, Table 2). GHG_c_ production was linear with time for all incubations. Average production rates and R^2^ values are included in Table 2. CO_2_:CH_4_ production ratios were significantly (p<0.05) elevated relative to median values between days 1-10—a phenomenon that we attributed primarily to increasing CH_4_ production rates. After this period, CO_2_:CH_4_ production ratios stabilized (as did CH_4_ production rates). We include only data collected after the stabilization period ended (day 10) in our calculations of average CO_2_:CH_4_ production ratios.

**Fig 2.**
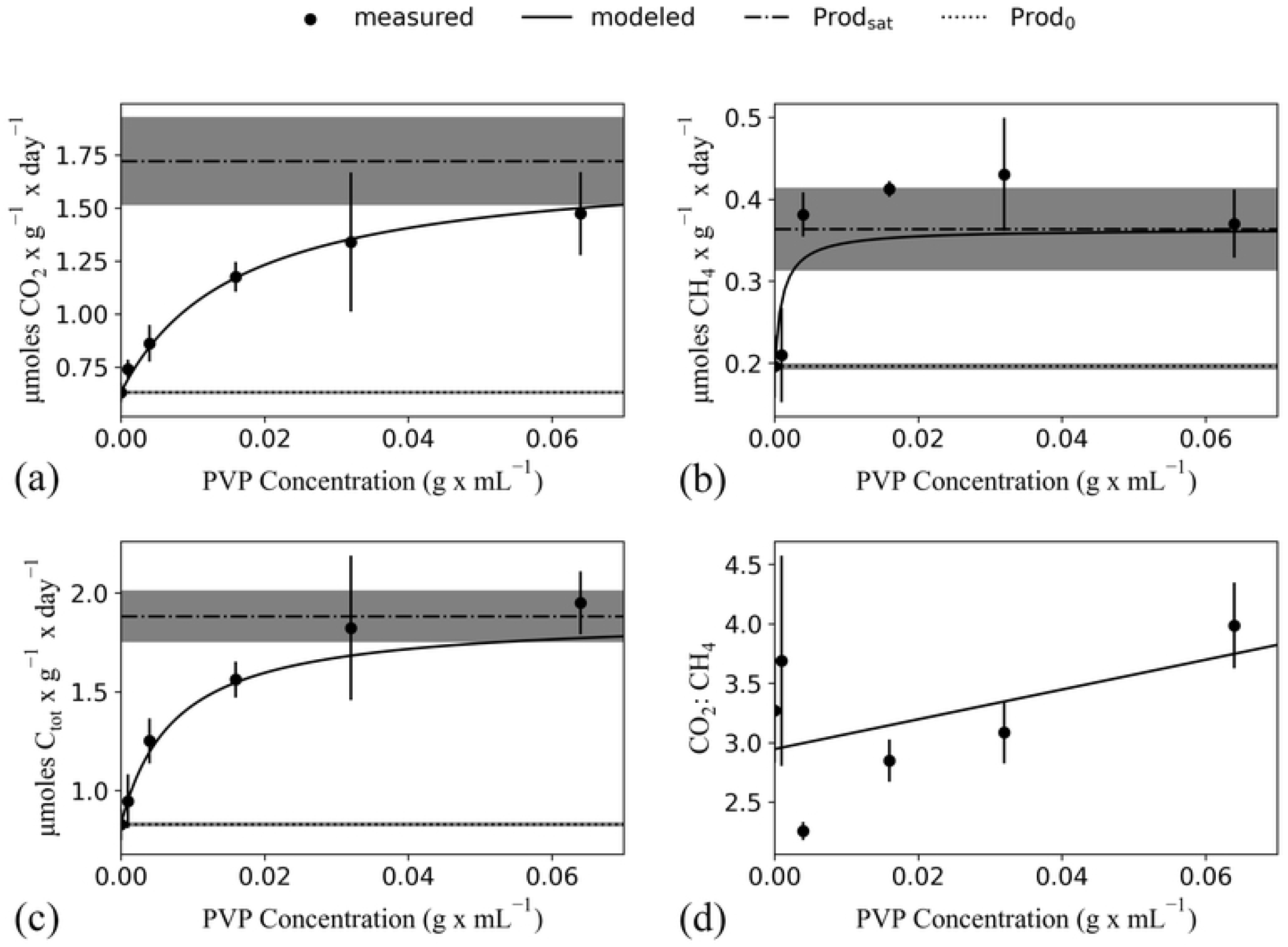
Observed and modeled impacts of polyvinylpyrrolidone (PVP) on C mineralization. PVP concentration (g mL^−1^) vs. (a) μmoles CO_2_ × g^−1^ × d^−1^ (R^2^ = 0.829, p< 0.001) (b) μmoles CH_4_ × g^−1^ × d^−1^ (R^2^ = 0.617, p< 0.001) (c) μmoles C_tot_ × g^−1^ × d^−1^ (R^2^ = 0.835, p< 0.001), and (d) CO_2_:CH_4_ (R^2^ = 0.414, p=0.65), where C_tot_=(CO_2_+CH_4_) and g=dry peat weight. Measured and modeled values are displayed as filled circles and solid lines, respectively. Modeled values were calculated using an amended Michaelis-Menten function (Eqn. 10, Methods) for panels a-c, and a best-fit linear regression curve for panel d. “Prod_sat_”—the estimated production rate for PVP-saturated peat (Eqn. 11, Methods)—and “Prod_0_”—the average control production rate—are displayed as dashed and dotted lines in panels a-c, respectively. Standard deviations for Prod_0_ and Prod_sat_ are depicted with gray shading.

**Table 2.** PVP concentration vs. production rate (C_tot_, CO_2_, and CH_4_) and average CO_2_:CH_4_.

| PVP Conc.<br>(g/mL) | $\text{CO}_2$ | | $\text{CH}_4$ | | $\text{C}_{\text{tot}}$ ($\text{CO}_2 + \text{CH}_4$) | | $\text{CO}_2:\text{CH}_4$ |
| --- | --- | --- | --- | --- | --- | --- | --- |
| | Prod. Rate | $R^2$ | Prod. Rate | $R^2$ | Prod Rate | $R^2$ | Average<sup>a</sup> |
| 0 | $0.63 \pm 0.04$ | 0.997 | $0.196 \pm 0.009$ | 0.964 | $0.83 \pm 0.04$ | 0.962 | $3.3 \pm 0.4$ |
| 0.001 | $0.74 \pm 0.06$ | 0.991 | $0.21 \pm 0.02$ | 0.949 | $0.95 \pm 0.08$ | 0.928 | $3.7 \pm 0.9$ |
| 0.004 | $0.86 \pm 0.06$ | 0.997 | $0.38 \pm 0.05$ | 0.985 | $1.3 \pm 0.1$ | 0.953 | $2.3 \pm 0.07$ |
| 0.016 | $1.18 \pm 0.08$ | 0.991 | $0.41 \pm 0.03$ | 0.981 | $1.56 \pm 0.08$ | 0.954 | $2.8 \pm 0.2$ |
| 0.032 | $1.34 \pm 0.08$ | 0.998 | $0.43 \pm 0.03$ | 0.992 | $1.8 \pm 0.4$ | 0.940 | $3.1 \pm 0.3$ |
| 0.064 | $1.5 \pm 0.1$ | 0.998 | $0.37 \pm 0.03$ | 0.966 | $2.0 \pm 0.3$ | 0.903 | $4.0 \pm 0.4$ |

Production rates ( $\mu\text{moles} \times \text{g}^{-1} \times \text{d}^{-1}$ ) were derived from best-fit linear regression lines.
<sup>a</sup>Average $\text{CO}_2:\text{CH}_4$ production ratios were calculated using values collected between day 10 and 57.

Amended Michaelis-Menten equations relating PVP concentration to GHG_c_ production (Eqn. 10, Methods) accounted for 83%, 62%, and 83% of the observed variance in CO_2_, CH_4_, and C_tot_ production, respectively (95% confidence interval, Fig 2, Michaelis-Menten parameters detailed in Table 3). PVP-saturated production rates (Prod_sat_; calculated using Eqn. 11, Methods) were significantly (p<0.05) higher than control production rates (Prod_0_; calculating by averaging control production rates), amounting to a 1.9, 2.3, and 2.4-fold increase in CO_2_, CH_4_, and C_tot_ production, respectively. CO_2_:CH_4_ production ratios ranged from 2.3-4.0 across all PVP concentrations studied. We approximated the relationship between CO_2_:CH_4_ vs. PVP concentration using a linear regression fit and found no significant correlation (R^2^=0.414, p=0.65; Fig 2) between these factors.

**Table 3.** Michaelis-Menten parameters for CO_2_, CH_4_, and C_tot_ (Fig 2a-c).

| Gas | k<sub>m</sub> | v<sub>m</sub> | Prod<sub>0</sub><sup>a</sup> | Prod<sub>sat</sub><sup>b</sup> | R<sup>2</sup> |
| --- | --- | --- | --- | --- | --- |
| CO<sub>2</sub> | 0.008 ± 0.004 | 0.08 ± 0.01 | 0.046 ± 0.004 | 0.13 ± 0.01 | 0.829 |
| CH<sub>4</sub> | 0.003 ± 0.002 | 0.02 ± 0.003 | 0.014 ± 0.003 | 0.037 ± 0.004 | 0.617 |
| C<sub>tot</sub> | 0.008 ± 0.003 | 0.11 ± 0.01 | 0.06 ± 0.006 | 0.17 ± 0.01 | 0.835 |

Michaelis-Menten equations relate PVP concentration (g × mL<sup>-1</sup>) to CO<sub>2</sub>, CH<sub>4</sub>, and C<sub>tot</sub> (CO<sub>2</sub>+CH<sub>4</sub>) production rate (in μmoles × g<sup>-1</sup> × d<sup>-1</sup>). k<sub>m</sub> and v<sub>max</sub> were calculated using the Python-based SciPy library with the Optimize package and the curve\_fit function. The standard deviation for these constants was determined via non-parametric bootstrapping (1,000 simulations). R<sup>2</sup> values reference the fit between modeled and measured values.
<sup>a</sup>Prod<sub>0</sub> (μmoles × g<sup>-1</sup> × d<sup>-1</sup>) refers to the average control production rate (where PVP=0 g · mL<sup>-1</sup>) and is equivalent to the y-intercept for the Michaelis-Menten curves.
<sup>b</sup>Prod<sub>sat</sub> (v<sub>max</sub>+Prod<sub>0</sub>) is an estimation of the production rate in PVP-saturated peat.

## Discussion

We hypothesized that increasing PVP concentration would yield increasing GHG_c_ production rates and that this relationship would follow an amended Michaelis-Menten (saturation) function (Fig 1). We aimed to quantitatively constrain the extent to which soluble phenolics inhibit C mineralization by cross-analyzing GHG_c_ production rates in control (Prod_0_) vs. PVP-saturated peat (Prod_sat_, calculated using Eqn. 11, Methods).

### Evaluating Hypotheses

The correlation between PVP concentration and GHG_c_ production was positive and robustly approximated using Michaelis-Menten curves (R^2^=0.829, 0.617, and 0.835 for CO_2_, CH_4_, and C_tot_, respectively). These findings are consistent with our hypothesis and can be reasonably attributed to PVP-phenolic interactions, which alleviate phenolic inhibition of microbial processes [24,25,43,44].

### Quantifying Phenolic Inhibition of C Mineralization

We calculated the extent to which phenolics apparently inhibited GHG_c_ production in our incubated bog peat using the following equation:

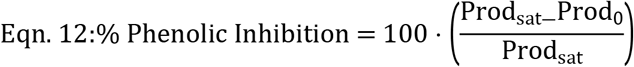

The reasoning behind this equation was as follows. PVP saturation is considered a method for decreasing soluble phenolic effectiveness. As such, GHG_c_ production in PVP-saturated peat (Prod_sat_) should theoretically approximate peat that is free of active soluble phenolics, except for the minute portion that persists in solution following equilibration with PVP. The %Phenolic Inhibition by phenolics should thus be measured by a comparison between PVP-saturated (Prod_sat_) and control (Prod_0_) peat. As it is feasible that persisting soluble phenolics could continue to react with enzymes [44], this equation offers a minimum estimate of %Phenolic Inhibition.

Using this equation, we determined that phenolics significantly (p<0.05) inhibited C_tot_, CO_2_, and CH_4_ production by 57 ± 16, 64 ± 24, and 46 ± 16%, respectively (Table 4; standard error calculating by propagating error from Prod_0_ and Prod_sat_).

**Table 4.** Percent Phenolic Inhibition of C_tot_, CO_2_, and CH_4_ Production Rates.

| Gas Analyzed | %Phenolic Inhibition<sup>a</sup> |
| --- | --- |
| C<sub>tot</sub><sup>b</sup> | 57 ± 16 |
| CO<sub>2</sub> | 64 ± 24 |
| CH<sub>4</sub> | 46 ± 16 |

<sup>a</sup>%Phenolic Inhibition estimates were calculated via eqn.12.
<sup>b</sup>C<sub>tot</sub> represents the sum of CO<sub>2</sub> and CH<sub>4</sub> production rates.

Quantifying differences between the soluble phenolic content in our incubations vs. field pore-water is beyond the scope of the study, which prevents us from applying %Phenolic Inhibition calculations directly to the field. To quantify the extent of this influence, it is necessary to identify potential differences between the abundance and speciation of soluble phenolics in incubation vs. field pore-water.

It is feasible that %Phenolic Inhibition in peat bogs varies significantly by site and depth, as associated differences in biotic (vegetation, humification indices, microbial community dynamics) and abiotic (temperature, pH) factors could collectively alter substrate supply and phenolic speciation. The former has been shown to significantly alter phenolic inhibition potential [9], while the latter remains largely unexplored. To broaden our assessments of %Phenolic Inhibition to wide geographic scales, further research investigating the interplay between phenolics and site/depth-specific parameters is needed.

If applicable to wide geographic scales, the strong inhibitory role of phenolics on C mineralization raises important implications. Numerous studies indicate that the introduction of molecular oxygen can stimulate phenol oxidase, which degrades phenolics, causing enhanced decay of organic matter [9,20,21,50]. This effect is thought to be exacerbated by (1) the onset of oxic respiration and (2) increased diversity and abundance of bacterial species capable of catabolizing phenolics—both of which have been observed following drainage of *Sphagnum*-rich peat [51]. These findings are the foundation of an assertion that rapid peat decay could occur following drainage (via anthropogenic disturbances) or exposure to drought (from rising temperatures) [9,20,21].

However, inconsequential effects of oxygen introduction on phenolic abundance have been observed in some peatland sites [36,52–54]. It has been suggested that peat bogs are particularly resilient to the effects of drainage, given that comparatively lower microbial community shifts following long-term drainage have been observed in bogs relative to fens [54]. Further research on this subject is needed before we can assess the implications of our %Phenolic Inhibition estimates on C mineralization shifts following water-table fluctuations.

If the findings presented herein apply to other permafrost-associated peatlands, habitat transitions due to thaw could significantly impact soluble phenolic content. Satellite studies have revealed widespread expansion of northern peatland shrubs over multidecadal timeframes [45,55], while warming experiments have induced significant loss of *Sphagnum* [56]. If the influence of phenolics on C mineralization is significantly higher in *Sphagnum*-rich (bog) vs. shrub-rich (fen) peatlands, the cumulative inhibition of peatland C mineralization by phenolics could be significantly offset by habitat transition.

Further research focusing on phenolic inhibition in fens is necessary to determine if this is the case. It is possible that the %Phenolic Inhibition of C mineralization in fens is similar to or even greater than bogs. If true, then the significant %inhibition estimates presented herein would not necessarily point to phenolics as significant contributors to the decreasing C mineralization rates typically observed across the fen-bog transition. On the other hand, significantly decreased %inhibition estimates in fens (relative to bogs) would suggest that phenolics do play pivotal roles in organic C retention within peat bogs.

## Conclusions

We employed an anaerobic incubation series of shallow water-logged bog peat (collected from Stordalen Mire, Sweden) inoculated with varying concentrations of PVP—a compound known to decrease phenolic effectiveness—as a means of investigating the impact of phenolics on C mineralization. We estimated that phenolics inhibit 57 ± 16% of total C mineralization (CO_2_+CH_4_) in the incubation conditions studied. This amount may underestimate the total inhibition since some phenolics may not be bound by PVP. To gain a full understanding of the phenolic impact on C mineralization throughout the subsurface peat column, further research identifying potential depth dependencies is necessary. To apply these findings globally, an understanding of regional controls on phenolic inhibition is necessary. Finally, cross-analyses between phenolic impacts on C mineralization in bog vs. fen habitats will allow us to determine whether the extraordinary recalcitrance in subsurface peat bog environments is attributed, at least partially, to phenolic activity.

## Acknowledgments

Funding for this study was provided by the Genomic Science Program of the United States Department of Energy Office of Biological and Environmental Research Grants (DE-SC0010580 & DE-SC0016440). Additional funding was provided by the EMERGE Biology Integration Institute of the National Science Foundation (NSF Award # 2022070). We want to thank collaborators from the department of Earth, Ocean, and Atmospheric Sciences at The Florida State University (FSU) for their assistance with the data analysis—especially Drs. Thomas Kelly and Michael Stukel. In addition, we gratefully acknowledge the staff of Abisko Naturvetenskapliga, and the 2018 IsoGenie field sampling team for collecting material used for the incubations.

